# Evidence of cryptic incidence in childhood diseases

**DOI:** 10.1101/079194

**Authors:** Christian E. Gunning, Matthew J. Ferrari, Erik Erhardt, Helen J. Wearing

## Abstract

Persistence and extinction are key processes in infectious disease dynamics that, due to incomplete reporting, are seldom directly observable. For fully-immunizing diseases, reporting probabilities can be readily estimated from demographic records and case reports. Yet reporting probabilities are not sufficient to unambiguously reconstruct disease incidence from case reports. Here, we focus on disease presence (i.e., marginal probability of non-zero incidence), which provides an upper bound on the marginal probability of disease extinction. We examine measles and pertussis in pre-vaccine era U.S. cities, and describe a conserved scaling relationship between population size, reporting probability, and observed presence (i.e., non-zero case reports). We use this relationship to estimate disease presence given perfect reporting, and define cryptic presence as the difference between estimated and observed presence. We estimate that, in early 20*^th^* century U.S. cities, pertussis presence was higher than measles presence across a range of population sizes, and that cryptic presence was common in small cities with imperfect reporting. While the methods employed here are specific to fully-immunizing diseases, our results suggest that cryptic incidence deserves careful attention, particularly in diseases with low case counts, poor reporting, and longer infectious periods.

## Introduction

### Epidemic Dynamics of Childhood Diseases

Measles and pertussis (whooping cough) are acutely infectious diseases caused by obligate human pathogens: the measles virus and *Bordetella pertussis*, respectively. These well-studied childhood diseases are fully immunizing but highly infectious, with a low average age of infection (< 10 years) in the pre-vaccine era [1]. Both diseases have fast life cycles compared to human host demographics [1].

Recurrent epidemics are a common feature of these diseases, driven by long-term host demographics and periodic forcing of disease transmission via changes in host density, such as school terms [2–4] or economic migration [5, 6]. At high incidence, susceptible hosts are rapidly depleted, leading to subsequent inter-epidemic troughs of low incidence, where stochastic extinction can occur. When infection is low or absent from a population, susceptible replenishment proceeds via the host demographic processes of birth and migration. These forces combine to yield characteristic yearly and multi-annual epidemic cycles in a range of diseases and human populations [7–14].

The life histories of measles and pertussis differ significantly in pace: measles has a shorter life cycle, is more “invasive”, and experiences more pronounced epidemics, while pertussis is the superior “colonizer”. The slower life history of pertussis is expected to dampen the effects of isolation relative to measles, and is predicted to enhance dynamical stochasticity [15, 16]. In pertussis, the contribution of waning immunity to observed dynamics has been a subject of extensive debate, both in infection-derived and vaccine-derived immunity [17–19]. In the pre-vaccine era, however, pertussis dynamics are consistent with the dynamics of infections that confer relatively long-lasting immunity, irrespective of whether the mechanism is long-term protection or natural immune boosting [20, 21].

Reporting probabilities vary widely between these diseases, as well as between locations [3, 22–24]. Measles infection causes characteristic symptoms of fever, rash, and pathognomonic Koplik’s spots [25]. Pertussis, on the other hand, exhibits age-dependent severity, and shares symptoms with many other common respiratory diseases [26, 27]. In addition, reports of pertussis in adults was generally absent in the pre-vaccine era [18]. Consequently, pertussis reporting is generally less complete and more variable than measles reporting [22, 24]. Such observational differences complicate meaningful comparisons between diseases, particularly in the presence of dynamical uncertainty.

### Determinants of Persistence

Persistence and stochastic extinction are key processes that affect pathogen ecology, evolution, and control efforts. As an ecological outcome, disease persistence arises from a complex interplay between local, within-population processes and metapopulation-level interactions among populations. Disentangling the impact of local and metapopulation processes on disease dynamics has proved challenging. At the local level, stochasticity in host and pathogen demographic processes commonly results in local extinction, particularly in small populations [28–31], and for pathogens with short infectious periods [31, 32]. Indeed, previous work has shown that local disease persistence scales approximately log-linearly with population size [31, 28, 33–35, 20, 36]. Likewise, theory predicts that, when all else is equal, longer latent and infectious periods and higher birth rates should increase local persistence [33, 31].

At the metapopulation level, host migration allows for imported infections to “rescue” local chains of infection [37, 36, 32, 38]. Intermediate levels of connectivity can aid rescue effects and metapopulation persistence [37], while very high levels of connectivity can synchronize populations and decrease rescue effects [39]. Low connectivity between populations, on the other hand, can favor boom-bust cycles. Here, disease importation is uncommon, and prolonged periods of local extinction allow susceptible individuals to accumulate far above equilibrium. Eventual pathogen re-introduction causes explosive epidemics that, in turn, reduce susceptible individuals far below equilibrium, thus favoring stochastic extinction.

Seasonal forcing plays a central role in these diseases, and can synchronize and/or accentuate periodic troughs of incidence in populations and metapopulations [11, 40, 41]. Yet the interplay between periodic forcing and temporal patterns of incidence can be be complex [42]. We note that previous work on measles has focused largely on England & Wales, and assumed a unified school calendar [2, 4, 40]. U.S. school calendars are set by local municipalities, and have varied considerably over both time and space [43–45]. As such, metapopulation patterns of extinction in response to seasonal forcing likely differ between these countries.

Here we focus on measles and pertussis in pre-vaccine era U.S. cities, where explosive epidemics and prolonged periods of low incidence and / or stochastic extinction are common. We examine a record of more than two decades of continuous weekly disease monitoring (1924-1945, Table 2) that includes the majority of U.S. urban areas in this era. The early 20*^th^* century U.S. provides an attractive model system: high-quality demographic records are available, and a diverse range of population sizes, ethnic compositions, and levels of geographic isolation are represented here. Life-long immunity provides a key dynamical constraint, allowing us to reliably estimate reporting probability. Finally, the absence of vaccination eliminates uncertainty associated with vaccine uptake and efficacy.

### Estimating Disease Presence

Due to imperfect reporting, the dynamical processes of persistence and stochastic extinction can seldom be directly observed. Previous work has estimated disease persistence from case reports, either in distinct human populations (e.g. cities, Conlan et al. [36]) or in metapopulations (e.g. countries, Metcalf et al. [32]). Lacking, however, are quantitative assessments of the impact of observational uncertainty on persistence estimates.

Species *presence* is a related quantity that has received considerable attention from community and conservation ecologists seeking reliable measures of species composition or richness. Here, sampling effort and species abundance have long been recognized to affect species detection probabilities [46, 47]. In assemblages of species, sampling effort can be accounted for via accumulation or rarefaction curves that quantify presence via asymptotic richness [47–49]. Related work has explored the interdependence between detection probability and spatiotemporal resolution [50], and has quantified the expected additional sampling required to achieve asymptotic detection [51].

Here we address a related problem: the reliable detection of a *single* species’ presence. In this case, reporting probability provides a proxy for sampling effort, while disease incidence is analogous to species abundance. We also explore the impact of temporal “grain size” [50] by aggregating case reports over a range of successively longer reporting windows. We suggest that the long-term, per-population probability of disease presence yields a lower bound on disease incidence, and provides an upper bound on time spent in an extinct state.

### Overview

We compare metapopulation patterns of presence between two diseases (measles and pertussis) within a single metapopulation. Disease incidence varies greatly over time, both within and across years; here we marginalize over time, and focus on long-term differences between populations and diseases. We use weekly, per-city disease case reports (*C_obs_*) and reporting probabilities to estimate the marginal (weekly) probability of disease presence (*P*). We show that city population size (*N*) and reporting rate (*r*) predict observed presence (non-zero case reports, *P_obs_*). We use this relationship to estimate the (weekly) probability of disease presence given full reporting (i.e. probability of non-zero incidence, *P_est_*). We find an increase in pertussis *P_est_* relative to that of measles across a range of population sizes. In addition, we show that the observed scaling of *P_obs_* with *N* and *r* is robust to temporal aggregation of case reports over longer reporting windows.

We define cryptic presence (*P_c_*) as the difference between estimated and observed presence: the (estimated, weekly) probability of unobserved presence. We show that *P_c_* scales with both population size and reporting probability, and is particularly common in small populations with low reporting probability.

## Methods

All subsequent analyses were conducted separately for each disease. All rates and probabilities are per week, unless otherwise noted. See Table 1 for definitions. Data are described in detail in Gunning et al. [24].

**Table 1:**
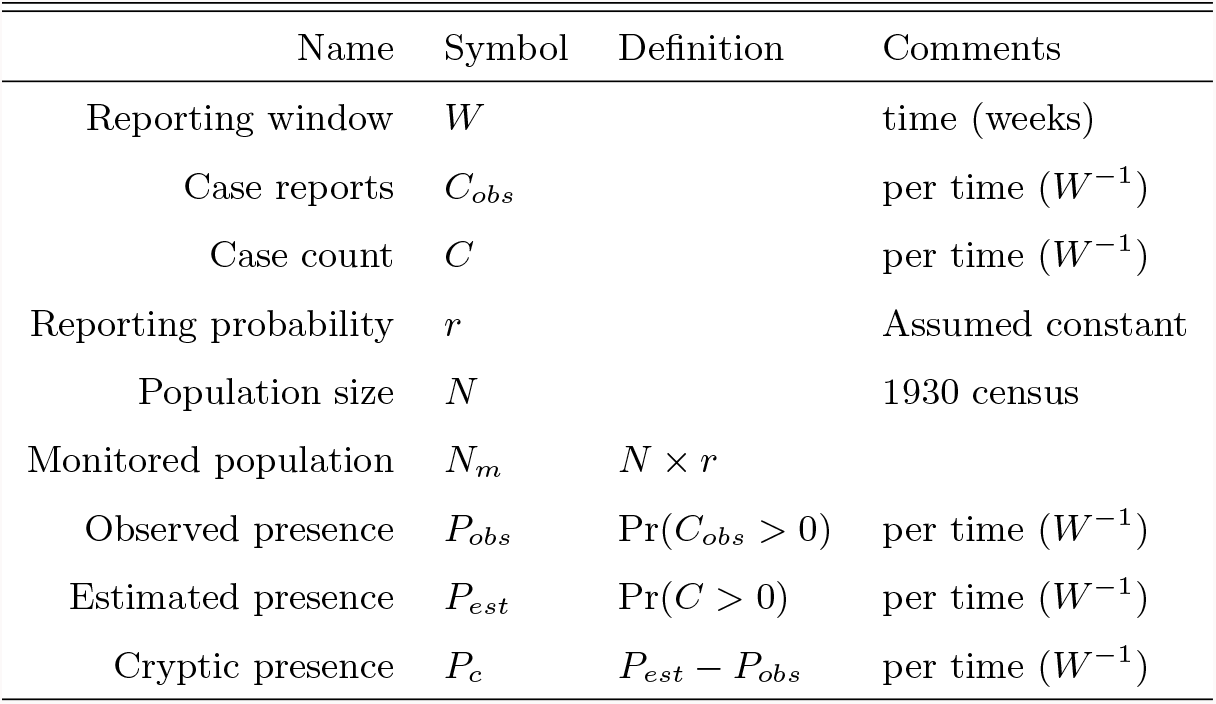
Definitions. Unless otherwise noted, a reporting window of one week was used (for case reports, case count, and presence probabilities).

### Incomplete observation

For each city and disease, we estimate a single, time-marginalized reporting probability (*r*) from case reports and demographic records, as in Gunning et al. [24]. We assume that each population’s proportion of susceptibles is in quasi-equilibrium over the period of record, and that the lifetime probability of infection is close to unity [3, 22]. As discussed in Gunning et al. [24], no strong evidence of time-variable reporting is apparent in this system over the period of record, and reporting probability is assumed to be constant over time.

We assume that case reports are generated via binomial sampling of cases: *C_obs_* ∼ Bin(*C, r*). Thus, each city’s *r* can be estimated from the ratio total case reports to total surviving births, summed over the period of observation: 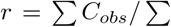 births. As described in Gunning et al. [24], total surviving births are estimated from yearly per capita state birth rates and infant mortality rates, along with yearly city populations. We also estimate approximate confidence intervals on *r* by bootstrapping (yearly) birth and infant mortality rates.

Given a binomial sampling process, we can estimate cases as the ratio of case reports to reporting probability: *C* = *C_obs_/r*. Yet this correction fails for *C_obs_* = 0, where our best estimate of *C* is zero. For low reporting probability and low incidence, a non-trivial proportion of observed zeros (i.e., *C_obs_* = 0) result from unobserved non-zero incidence. Consider, for example, *r* = 0.1 and *C* = 10, such that Pr(*C_obs_* = 0) = (1 *− r*)*^C^* ≈ 0.35. Here, approximately 35% of case reports will be zero, thus yielding erroneous under-estimates of *C* = 0. In short, observed zeros result from a “mixed process” of disease absence (*C* = 0), together with unobserved, cryptic incidence: (*C* > 0 ∩ *C_obs_* = 0). It is this unobserved presence of disease that we seek to quantify.

### Estimated and Cryptic Presence

As noted above, we focus here on the marginal, per time probability of disease presence. We first exclude cities where a disease was always present (Pr(*C_obs_* > 0) = 1). We define the monitored population (*N_m_*) as the full population scaled by the reporting probability: *N_m_* = *N × r*. Note that, at full reporting, *N* = *N_m_*.

We employ a binomial generalized linear model (B-GLM) to model the response of *P_obs_* to log *M_m_*, where disease presence (*C_obs_* > 0) is equated with the binomial trial’s “success”. Reporting weeks with NAs were excluded, and each city was weighted by the number of non-excluded reporting weeks. We estimate a single 2-coefficient B-GLM (slope + intercept) for each disease using R’s glm interface [52].

We use a complementary log-log (cloglog) link function (*f*): *f* (*P_obs_*) ∼ log *N_m_*. Unlike the more common logit link, the cloglog link hypothesizes an asymmetric response to the predictor. That is, at large population sizes, cities approach complete presence (*P_obs_* = 1) more rapidly than a logit link predicts. This accords with biological intuition, where mechanistically distinct processes dominate near complete presence versus near complete absence. Further discussion of mechanistic biological interpretations of this model formulation is included below.

We use the resulting B-GLMs to extrapolate estimated disease presence from full population size: *f* (*P_est_*) ∼ log *N*. As noted, *N* simply equals the monitored population under complete reporting, motivating our choice.

Estimated presence is the sum of observed presence and cryptic (unobserved) presence (*P_c_*): *P_est_* = *P_obs_* + *P_c_*. As such, we estimate cryptic presence as the difference between estimated and observed presence: *P_c_* = *P_est_ − P_obs_*. Non-zero cryptic presence, in turn, provides evidence of cryptic incidence.

### Model Exploration

We also explore the effect of the length of reporting windows. We sum case reports over reporting windows of varying widths *W*, ranging from 2 to 16 weeks. To compute reporting window sums (*C_obs,W_*), NA weeks were omitted, and windows containing only NA weeks were excluded (excluded windows were common for pertussis). *P_obs,W_* was computed as the proportion of non-zero window sums: Pr(*C_obs,W_* > 0). We then build a new model for each disease using both log *N_m_* and *W* as predictors of *C_obs,W_*.

The cloglog link function *f* (*x*) = log(*−* log(1 *− x*)) also provides a biologically relevant hazard analysis interpretation of the postulated model formulation. For each population, assume a constant rate of infection (*λ*) and total susceptible population (*S*). Then the probability of no new infections in a given time window *W* is Pr(*C* = 0) = 1 *− P ≈* exp(*−λSW*). For a given population size *N*, the susceptible proportion is then *S/N*, and *P* = Pr(*C* > 0) = 1*−*exp(*−λN* (*S/N)W*). The cloglog link then yields: *f* (*P*) = log(*λ*)+log(*S/N*)+log(*N*)+log(*W*). Thus, we expect the transformed response (*f* (*P*)) to change linearly in both log(*N*) and log(*W*). In truth, the populations studied here are not at equilibrium, such that *λ* and *S/N* instead oscillate over time around a long-term mean. Nonetheless, the above analysis hypothesizes a functional relationship between *P*, *N*, and *W*, which we explore further below.

### Estimating uncertainty

Both *P_est_* and *P_c_* are influenced by uncertainty arising from estimates of *r*, as well as GLM predictions. For each disease, we used a two-step process of bootstrap resampling to estimate the combined impact of reporting probability and B-GLM prediction uncertainty. Note that population size *N* changes over the period of record, and no uncertainty or variation therein is accounted for here.

First, bootstrap draws of *r* (henceforth *r_b_*) were taken via non-parametric resampling. For each draw, yearly state birth rates and national infant mortality rates were resampled, and total births thus summed (see Gunning et al. [24] for details). The resulting *r_b_* were used to compute *N_m_* from *N*, and a B-GLM was fit to the result. In this way, 1e + 04 models were fit.

These models were then used to extrapolate *P_est_* from *N*. The following was conducted for each bootstrap model (above), and for each city within that model. To incorporate per-city variance of *r* into model predictions, *N* was back-estimated from a (new) bootstrapped *N_m_*, which was then divided by the estimated reporting rate: *N* = *N ×* (*r_b_/r*). The model’s expected value of *P_est_* was then extrapolated from *N*. A random binomial sample was then taken, where the number of trials equaled the number of non-NA weeks for that city, with Pr(*success*) = *E*(*Pest*). The bootstrap draw of *P_est_* is then the proportion of successes.

Finally, *P_c_* was computed from *P_obs_* and the re-sampled *P_est_*. The resulting bootstrap samples were used to construct prediction intervals (95% PI) for *P_est_* and *P_c_*.

## Results

For reference, time series of sum case reports, as well as variance-scaled case reports of select cities, are shown in Figures S2 and S3. Summary statistics and observation counts are shown in Table 2. Overall, the average reporting probability of pertussis is much lower than for measles, with a higher coefficient of variation among cities, as discussed in Gunning et al. [24]. Figure 1 shows *P_obs_*, *P_est_*, and *P_c_* (rows) versus *N* and *N_m_* (columns). Figure 1 also provides a visual illustration of *P_est_ − P_obs_* = *P_c_*, i.e., panels *E* = *C − A*, and *F* = *D − B*.

**Figure 1:**
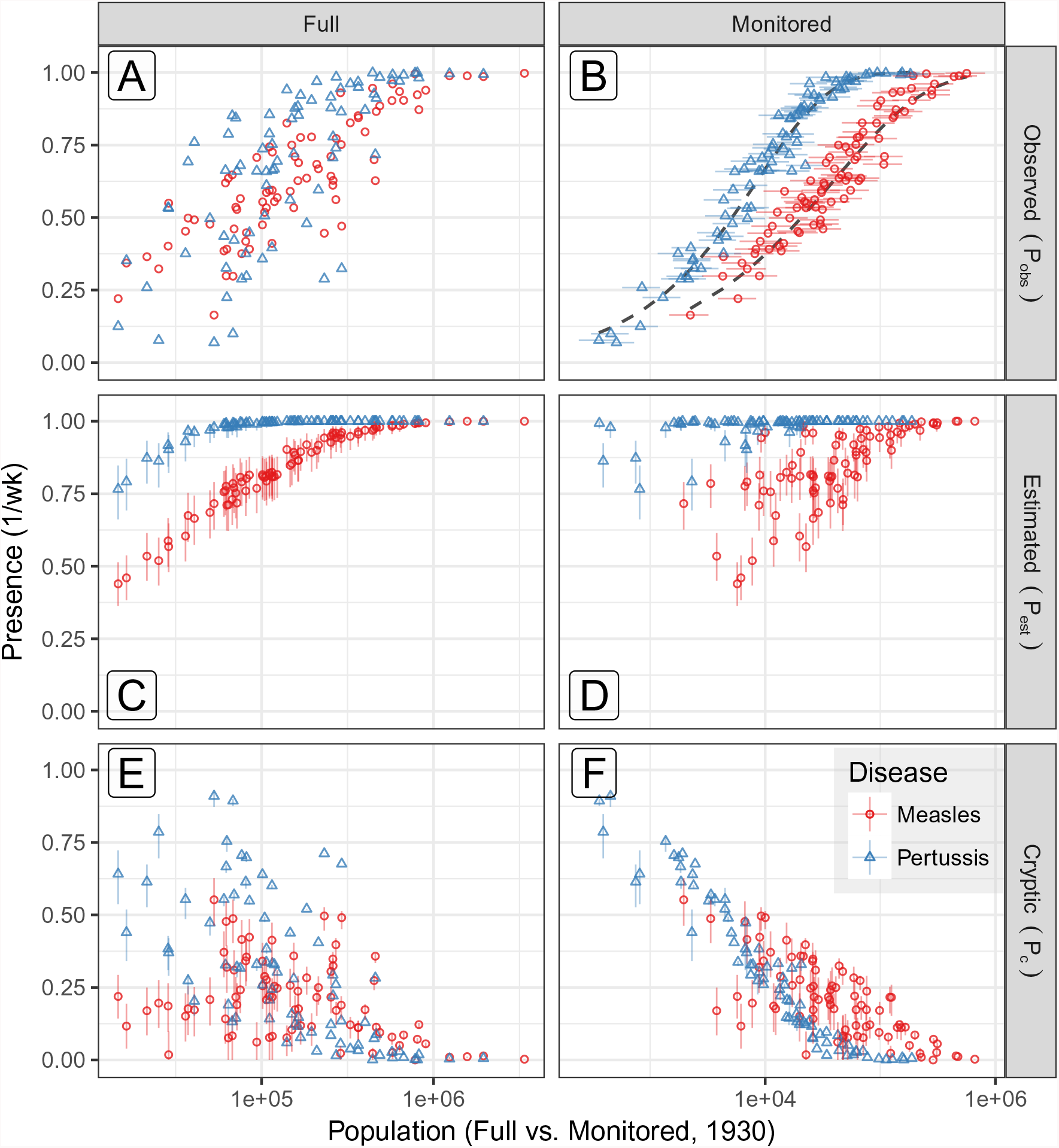
Presence (*P*) by population size (*N*). Columns: full population (*N*) and monitored population (*N_m_* = *N × r*). Rows: observed (*P_obs_*), estimated (*P_est_*), and cryptic (*P_c_* = *P_est_ − P_obs_*) presence. **A:** Empirical observations of *P_obs_* versus *N*. **B:** *N* is scaled by incomplete reporting to yield *N_m_*. Horizontal bars show uncertainty in *r* (95% CI). The response of *P_obs_* to *N_m_* is modeled with a binomial GLM (cloglog link, one model per disease). Dashed black lines show model fits (see Figure S1 for details). **C:** The resulting models are used to predict disease presence at full reporting (*P_est_|r* = 1), along with 95% PI. Here, *P_est_* of pertussis is higher than measles in all but the largest cities. **D:** As in C, but with *N_m_* (see below). **E,F:** For each city, cryptic presence (*P_c_*) is the difference between the previous two rows: *E* = *C − A*, and *F* = *D − B*. **E:** *P_c_* is uncommon in large cities, likely due to the larger number of total cases per week. Panel **F** shows that, for pertussis, *P_c_* increases predictably with both population size and reporting. Measles, on the other hand, shows considerable variation in the response of *P_c_* to *N_m_*, suggesting a non-linear response of disease incidence to city size.

**Table 2:**
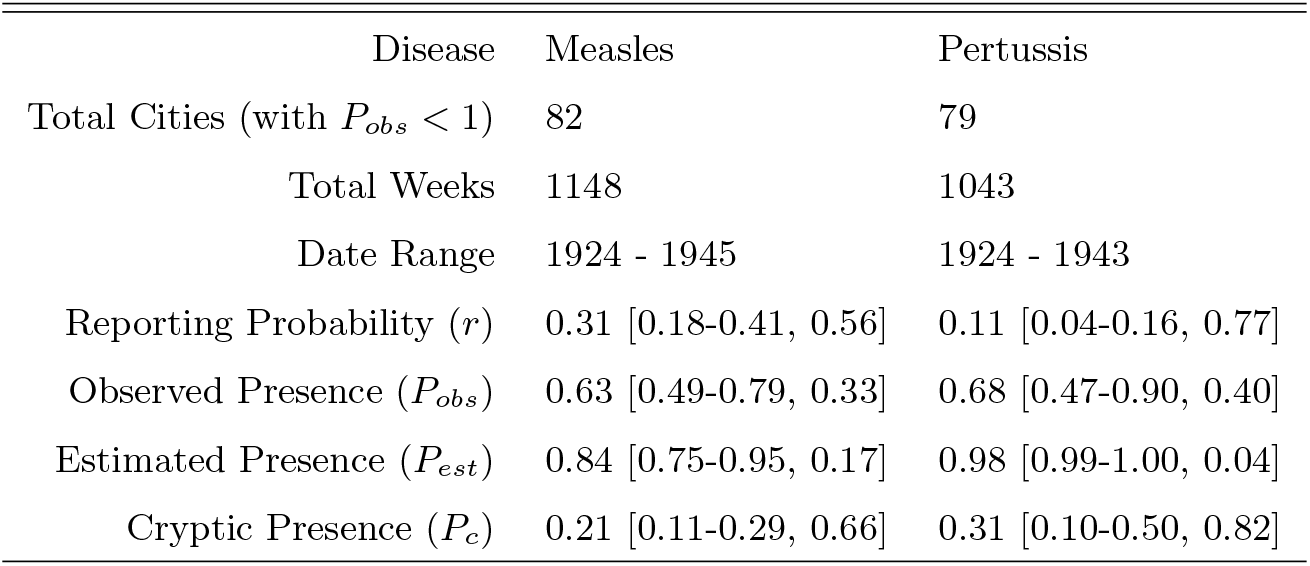
Overview: number of included cities, time period of record (inclusive), and summary statistics for studied cities: (Mean [Q1-Q3, CV]), including observed, estimated, and cryptic presence (*P_obs_*, *P_est_*, and *P_c_*, resp.).

Regardless of disease, we expect a lower probability of presence in smaller populations [28–31]. Indeed, we find that population size (*N*) predicts observed presence (*P_obs_*). As shown in Figure 1A, no difference between diseases is evident when solely case reports are considered. When reporting is considered, however, monitored population size (*N_m_*) yields an excellent predictor of *P_obs_* (Figure 1B): pseudo-*R*^2^ = 0.908 (measles) and 0.958 (pertussis). We use the models shown in Figure 1A to predict *P_est_* from *N*. The resulting predictions are shown in Figure 1C (plotted against *N*) and Figure 1D (plotted against *N_m_*). Cryptic persistence (*P_c_*) is simply *P_est_ − P_obs_*, such that Figure 1E is the difference between Figure 1C and Figure 1A.

Theory predicts that pertussis, with a longer infectious period and lower transmission rate, should exhibit less frequent stochastic extinction than measles for a given population size [33, 31], a pattern obscured by pertussis’ low and variable reporting. Correcting for incomplete reporting, we estimate that pertussis presence (*P_est_*) is indeed higher across a wide range of population sizes (Figure 1C).

As expected, cryptic presence of both diseases is rare in large populations (Figure 1E), where case count is high. In the remaining populations, however, the two diseases differ. For measles, cryptic presence is common across a wide range of population sizes, though true absence (i.e. via stochastic extinction) appears to dominate in smaller cities (Figure 1C). For pertussis, cryptic presence is most common in smaller cities, where frequent failures to detect disease arise from a combination of low reporting and low case count (Table 2, Figure 1E).

We expect cryptic presence to be a function of both reporting probability and the underlying distribution of case counts. Indeed, we observe increasing *P_c_* with decreasing *r* for both diseases (Figure 2). We also find marked differences between diseases: for a given *r*, measles generally experiences higher *P_c_*, possibly due to prolonged periods of low incidence. Finally, conditioned on *r*, larger populations exhibit lower *P_c_* than smaller populations, particularly for pertussis (Figure 2, inset).

**Figure 2:**
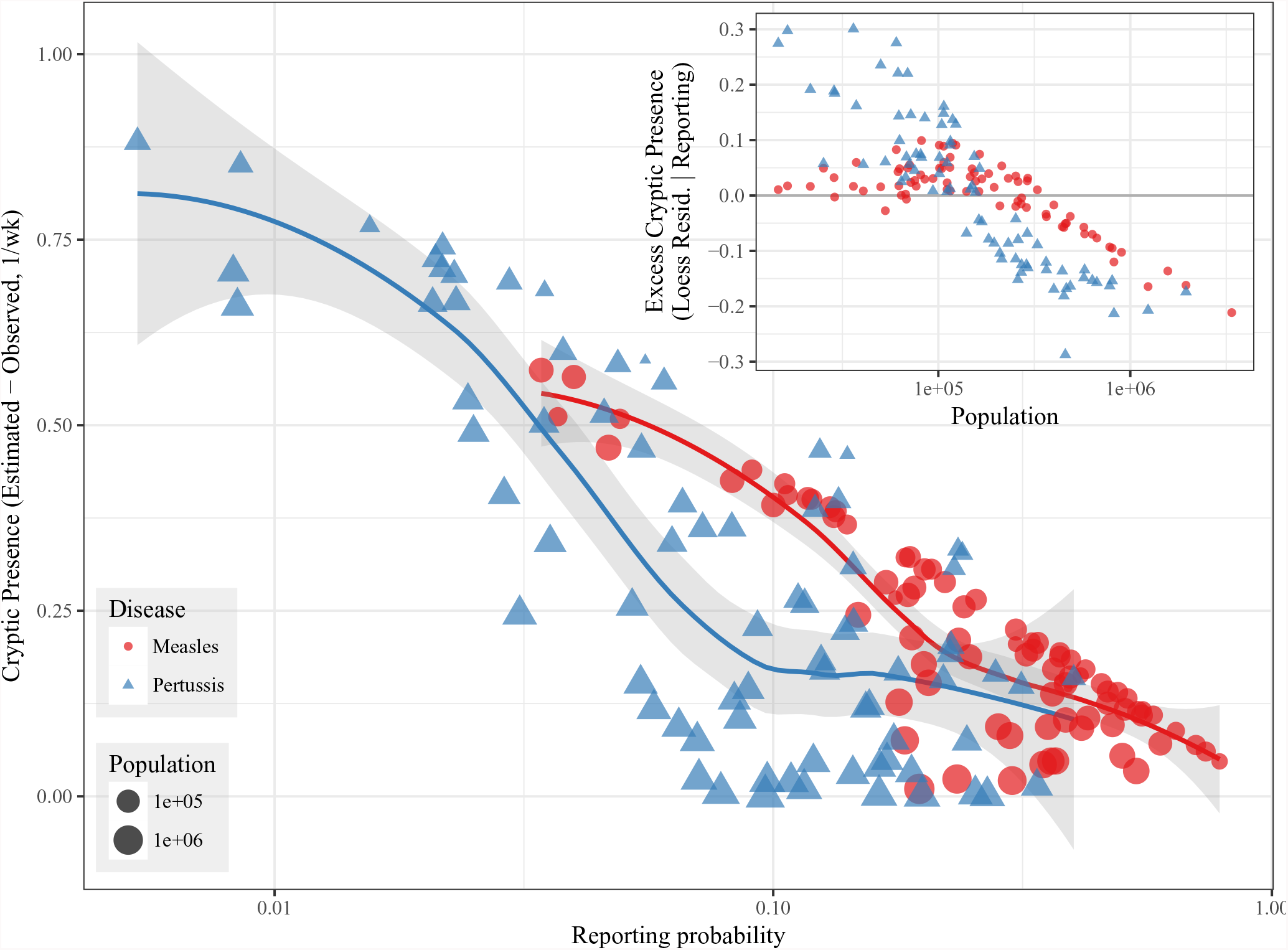
Cryptic presence (*P_c_*) by reporting probability (*r*). Reporting of pertussis is less complete and more variable than measles; cryptic presence also varies widely in pertussis. A superimposed LOESS regression shows that, at low reporting probabilities, cryptic presence is strongly correlated with reporting probability. The residuals of the LOESS regression are also plotted against population size (inset figure). For a given reporting probability, larger cities generally exhibit less cryptic presence than smaller cities, particularly for pertussis. Cryptic presence is essentially absent in the largest cities, regardless of disease or reporting probability.

### Temporal aggregation

Model fits, including the effects of temporal aggregation, are shown in Figure S1 and Table S1. As predicted, we find that (cloglog-transformed) *P_est_* increases linearly in both log(*N*) and log(*W*). The slope of *P_est_* in response to *N* is steeper in pertussis, suggesting that pertussis reaches complete presence more quickly than measles with increasing population size (as theory predicts).

Table S1 also shows the gradual decay of model fidelity with increasing temporal aggregation, along with an associated reduction in sample counts, as cities with complete presence (*P_obs_* = 1) are omitted. A close inspection of Figure S1A also reveals, at high levels of aggregation, poor model fits in large populations, where observed presence is far below model predictions (i.e. large negative residuals). This pattern likely results from the small number of available reporting windows, limiting the range of values that *P_obs_* can adopt. At *W* = 16 weeks, for example, the maximum number of (non-excluded) reporting windows per city is 61 (pertussis) and 71 (measles), such that the maximum incomplete *P_obs_* is approximately 0.984 and 0.986, respectively.

## Discussion

Despite widespread availability of inexpensive and effective vaccines, childhood diseases have resisted elimination efforts. Classic epidemiological theory proposes that reducing the susceptible proportion of a population below 1*/R*0 should interrupt disease transmission, leading to local extinction [1]. Yet metapopulation elimination of disease has proven elusive and expensive: morbidity and mortality from vaccine-preventable diseases remains high in developing nations [53, 54], and importation of infection back into previously disease-free populations and metapopulations continues [55–57].

Where, when, and why vaccine-preventable diseases persist are key ecological questions with important modern epidemiological consequences. As we have shown, incomplete disease reporting substantially affects common measures of disease presence, particularly for low reporting probability and low case counts. This impedes inference about disease dynamics at the local scale, and complicates comparisons between diseases or metapopulations with different reporting probabilities.

One particular area of practical concern that warrants increased attention is the fidelity of available demographic records in the modern era. Birth and migration rates help constrain reporting estimates and inform control measures [58]. Unfortunately, low birth registration coverage is common in modern developing nations [59], where incidence of vaccine-preventable diseases such as measles is currently highest [60]. In addition, completeness of birth registration varies greatly by geographic region and socioeconomic status [59, 61]. In some cases, multiple independent sources of demographic records can be employed to validate findings, such as the use of both government census records and survey-based Demographic and Health Surveys [62].

A key challenge in disease ecology is the unraveling of complex feedbacks between metapopulations and their constituent populations. Local disease persistence is driven both by local processes (birth, disease transmission) and metapopulation processes (host migration, disease importation). This study system pairs two different diseases within the same metapopulation, highlighting differences due to pathogen life history.

Here we estimate that cryptic presence is widespread in both diseases. We expect that cryptic presence is concentrated in cities that exhibit long periods of low but non-zero incidence, teetering on the edge of stochastic extinction. Yet the characteristics of these “refuge” populations differ markedly between diseases. We find that cryptic presence is concentrated at smaller populations in pertussis than in measles (Figures 1E and 2). This accords with epidemiological theory, which predicts that measles’ high transmission rate and short infectious period leads to rapid susceptible depletion in small populations. Thus, small populations are expected to commonly experience measles extinction. Pertussis, on the other hand, can sustain low but non-zero incidence in much smaller populations than measles due to a longer infectious period and lower transmission rate.

### Relation to Previous Work

Incomplete reporting is a common feature in human diseases, but has received relatively little attention. A comprehensive review of incomplete reporting in this system is given in Gunning et al. [24]. Recent analysis of historical U.S. polio concludes that “absence of clinical disease is not a reliable indicator of polio transmission” due to unobserved incidence [63], highlighting the critical role incomplete reporting can play in modern disease control.

Critical community size is one commonly employed threshold measure of disease persistence. CCS in particular, and threshold measures of extinction in general, has been widely criticized as poorly specified and difficult to measure [31, 64, 38]. In addition, cryptic presence should artificially inflate CCS estimates, as larger populations appear to undergo stochastic extinction. Nonetheless, the CCS of a disease remains a commonly reported “feature” of empirical data. For comparison, we present a simple empirical definition of CCS: the minimum population size where observed or estimated presence (*P_obs_* and *P_est_*, resp.) exceeds 95% (i.e., min(Population) given *P* > 0.95; Figure 3). The effects of incomplete reporting here are dramatic: for measles, CCS changes from ≈ 580 thousand (*P_obs_*) to ≈ 330 thousand (*P_est_*), while for pertussis, CCS changes from ≈ 210 thousand (*P_obs_*) to 50 thousand (*P_est_*).

**Figure 3:**
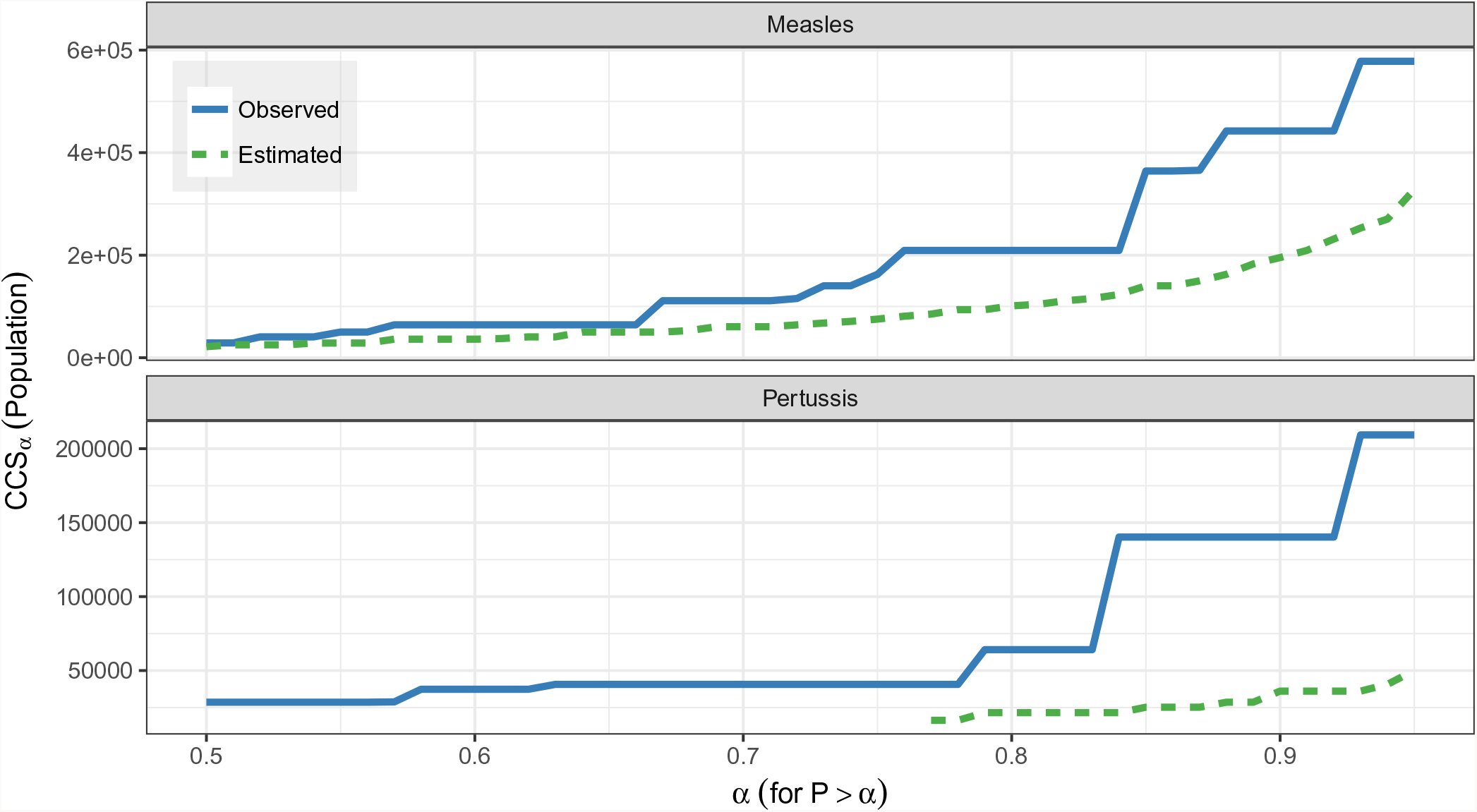
Empirical estimates of *CCS_α_*: the minimum population size (*N*) such that *α < P* (for 0 *< α <* 1). Results shown for both observed presence (*P_obs_*, blue dashed line) and estimated presence (*P_est_*, green solid line). Thus, *CCS_α_* is the minimum *N* where the disease is present more than *α* proportion of sampled weeks. Pertussis is estimated to be present in all cities at *α* = 0.76 (i.e., present in more than 76% of sampled weeks).

We expect that lower metapopulation incidence should, in general, decrease local persistence by reducing disease importation. How local persistence scales up to metapopulation persistence is less clear. Conventional epidemiological wisdom [65, 41] holds that metapopulation persistence depends on local persistence in focal cities above a critical size (i.e., CCS). Recent work suggests that aggregates of medium-sized cities exhibit patterns of persistence similar to individual cities of comparable size [38]. Our estimates of widespread cryptic presence in cities experiencing low case counts further emphasizes the role that “non-focal” cities can play in metapopulation persistence.

### Implications for Disease Detection and Control

Recent detection of wild-type polio in Nigeria [66] clearly illustrates that failure to account for cryptic presence can lead to biased assessments of control effort efficacy, and mistaken allocation of control efforts away from areas where disease remains present. Previous work has demonstrated the high likelihood of unobserved incidence in polio, where low case counts and poor reporting commonly co-occur [63]. Our results provide a clear warning against overly optimistic interpretations of apparent disease absence, and the critical importance of ongoing surveillance efforts.

More generally, the observed interdependence between cryptic presence, incomplete reporting, and case counts adds uncertainty to ongoing disease control efforts. As case counts drop, the frequency of cryptic presence is expected to become more sensitive to incomplete reporting. On the other hand, successful control measures will potentially lower cryptic presence in small populations, as those populations transition from low but non-zero incidence into true extinction. Indeed, this pattern has been observed in pertussis in England & Wales [42].

Here we show that cryptic presence can have a complex and disease-specific relationship with population size; we also provide a method for estimating this relationship from historical surveillance records in fully immunizing diseases such as polio or rubella. While these *methods* are not directly applicable to multi-strain diseases such as influenza or dengue, or repeat infections such as malaria, we suggest that a similar relationship between incomplete monitoring and undetected incidence is likely, based purely on intrinsic sampling stochasticity in real-world disease detection.

In practice, these methods could assist in the optimal allocation of resources for an active surveillance strategy of a fully-immunizing disease: first, to identify when elimination has been achieved at the meta-population scale and second, to monitor the maintenance of elimination. The results presented here suggest, for example, that additional resources for pertussis monitoring would be best allocated towards active surveillance in smaller populations (Figure 1E).

An additional complication is that disease monitoring intensity is commonly tied to disease incidence. This could lead to the paradoxical increase in cryptic presence as a disease approaches elimination due to reduced monitoring efforts. One example is pertussis, where high vaccination rates in developed nations have decreased incidence to very low levels [67, 42, 68]. In some locations, low incidence has led to the cessation of routine disease surveillance [26]. Active surveillance, on the other hand, has revealed widespread unreported incidence [26], including asymptomatic infection and subsequent transmission [69, 70].

For immunizing diseases, cryptic incidence does serve to increase natural immune boosting, even in the presence of widespread vaccination. The well-known “honeymoon period” [71] refers to the combined benefits of disease-induced and vaccine-induced immunity in a population shortly after the introduction of vaccination. As disease incidence falls, however, disease-induced immunity drops, leading to paradoxical negative feedback between vaccine-induced immunity and immunity from natural infection [58]. Cryptic incidence again adds an element of uncertainty regarding the long-term immune status of populations. Our results suggest that active monitoring could be used to identify sero-conversion or immune boosting [27, 21] from cryptic incidence, which could, in turn, inform ongoing control efforts.

The above-noted uncertainties highlight the need for novel, cost-effective monitoring to assess the frequency of cryptic presence. One example is genetic sequence monitoring, which could provide evidence of local or metapopulation persistence. Increased awareness of pathogen persistence could, in turn, inform phylodynamic models that seek to couple the ecology and evolution of human diseases [72–74], as well as provide novel insight into patterns of host metapopulation connectivity.

Here we use a well-studied, highly-constrained system to show that cryptic presence was both common and explicable in the pre-vaccine era U.S. Our work, along with recent public health developments, suggests that attention to cryptic presence in other disease systems is warranted. Widespread asymptomatic malaria incidence in Southeast Asia, for example, has been suggested as a potential reservoir of artemisinin-resistant *Plasmodium falciparum* [75, 76], whose spread represents a “major threat to global public health” [77]. In the case of wild-type polio, eradication of the last few cases has proved both expensive and logistically challenging [78]. After two years of apparent disease absence, the recent detection wild-type polio virus’ endemic persistence in Nigeria argues strongly against complacency in disease surveillance efforts [66]. An improved understanding of the relationship between disease monitoring effort and cryptic disease presence can benefit modern and future disease control efforts.

## Author Contributions

CEG and HJW designed the study. CEG performed the analyses and wrote the first manuscript draft. EBE contributed to design and execution of analyses. All authors contributed to subsequent manuscript revisions.

## Acknowledgments

Comments from four anonymous reviewers greatly improved the manuscript.

The authors would like to thank Natalie Wright, Michael A. Robert, Michael Chang, James H. Brown, and Melanie E. Moses for their support and assistance.

MF was supported under a grant from the Bill and Melinda Gates Foundation and the RAPIDD Program of the Science and Technology Directorate of the Department of Homeland Security. CG was supported by a fellowship in the Program in Interdisciplinary Biological and Biomedical Sciences at the University of New Mexico. This publication was made possible by Grant Numbers P20RR018754 from the National Center for Research Resources (NCRR), T32EB009414 from the National Institute of Biomedical Imaging and Bioengineering (NIBIB), components of the National Institutes of Health (NIH). Its contents are solely the responsibility of the authors and do not necessarily represent the official views of NCRR, NIBIB, or NIH.

## Data Accessibility

- U.S. case report data and demographics available at Data Dryad, doi:10.5061/dryad.92p46

## Supplemental Information

**Table S1:** **Binomial GLMs predict** the response of observed presence (*P_obs_*) to monitored population size (log *N_m_*), for different temporal aggregation lengths (*W*, weeks). To aggregate case reports, missing weeks are omitted, and reporting windows containing only missing weeks are excluded. For each *W* and city, 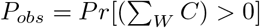. Cities with *P_obs_* = 1 (disease always present) are omitted, and cities are weighted by the number of non-excluded reporting windows. As aggregation increases, the number of available cities decreases, as does the number of reporting windows. Pseudo-*R*_2_s (proportion explained deviance) show a decrease in *R*_2_ at large *W* (see Figure S1). For illustrative purposes, each table row shows a separate model (i.e., each disease and *W*).

| Disease | Window Length ($W$, weeks) | # of Cities | # of Windows | Pseudo-$R^2$ |
| --- | --- | --- | --- | --- |
| Measles | 1 | 82 | 1147 | 0.908 |
| Measles | 2 | 80 | 573 | 0.886 |
| Measles | 4 | 74 | 286 | 0.827 |
| Measles | 8 | 59 | 143 | 0.715 |
| Measles | 16 | 34 | 71 | 0.635 |
| Pertussis | 1 | 79 | 1042 | 0.958 |
| Pertussis | 2 | 73 | 521 | 0.937 |
| Pertussis | 4 | 59 | 259 | 0.888 |
| Pertussis | 8 | 46 | 130 | 0.848 |
| Pertussis | 16 | 32 | 61 | 0.732 |

**Figure S1:**
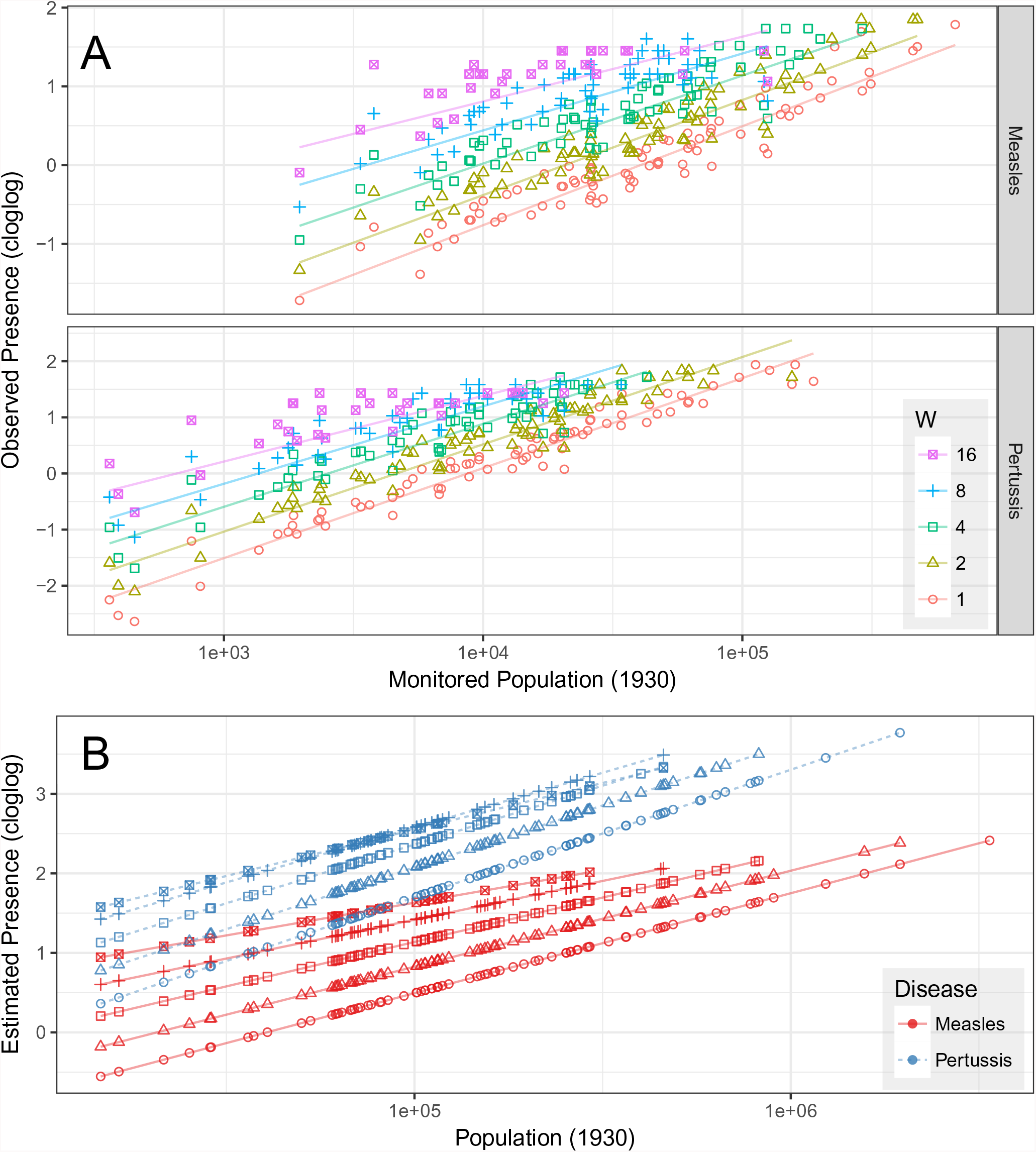
Binomial GLMs: **A: Observed Presence.** Response of observed presence (*P_obs_*) to monitored population size (*Nm*) and reporting window length (*W*, weeks). Lines show model estimates, with a separate model for each disease, using a cloglog link (y-axis, *f*): *f* (*P_obs_*) ∼ log *Nm|* log *W*. See Table S1 for details and observation counts. **B: Estimated Presence.** Model estimated presence (*P_est_*) in response to population size (*N*, 1930): *f* (*P_est_*) ∼ log *N |* log *W*.

**Figure S2:**
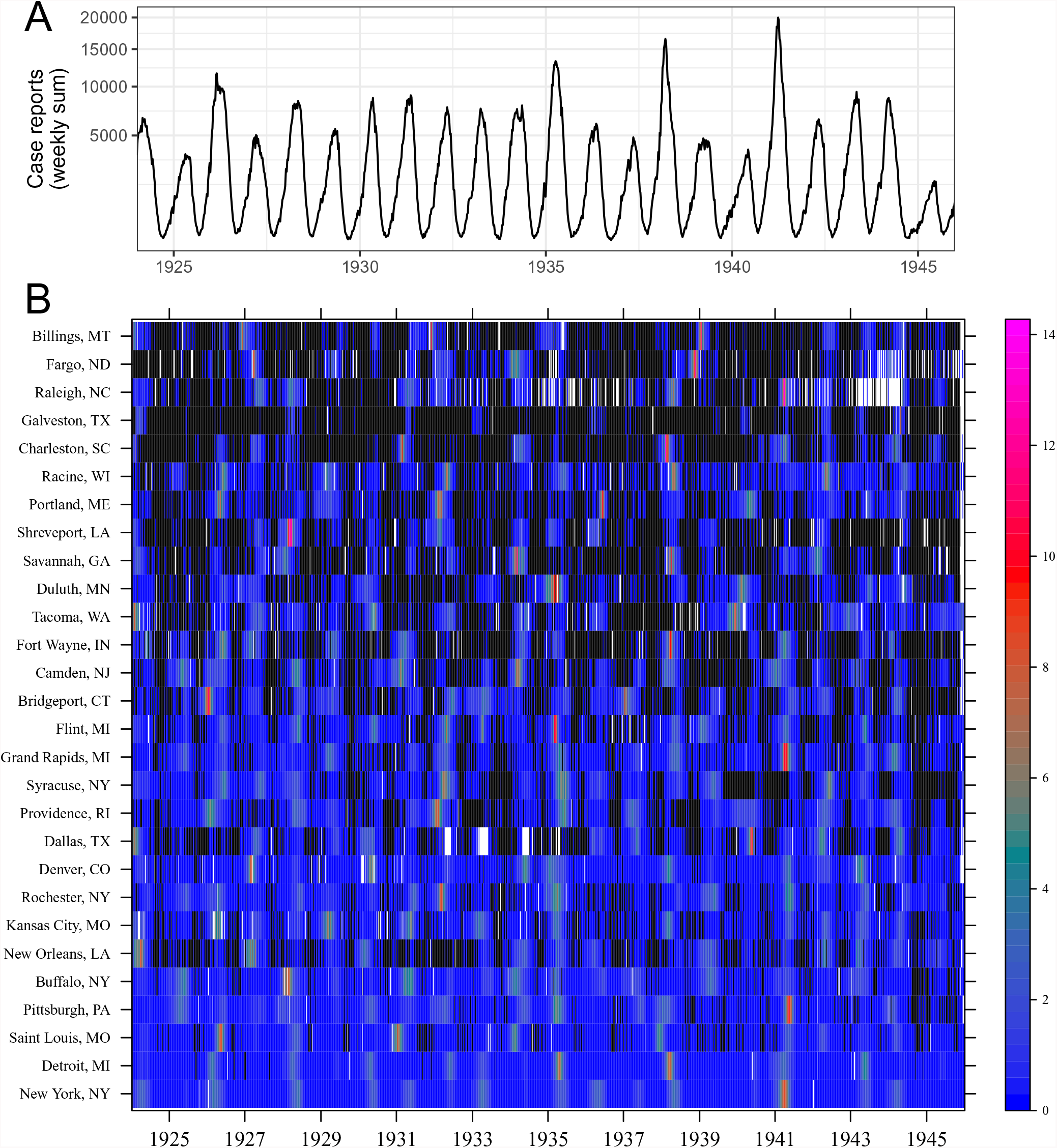
**Measles. A**. Total weekly case reports (all cities, sqrt-transformed). **B**. Weekly case reports (black = 0; white = missing). One city per row, ordered by population size, showing every third city. To facilitate comparison between cities, each city’s observations are scaled by that city’s temporal variance.

**Figure S3:**
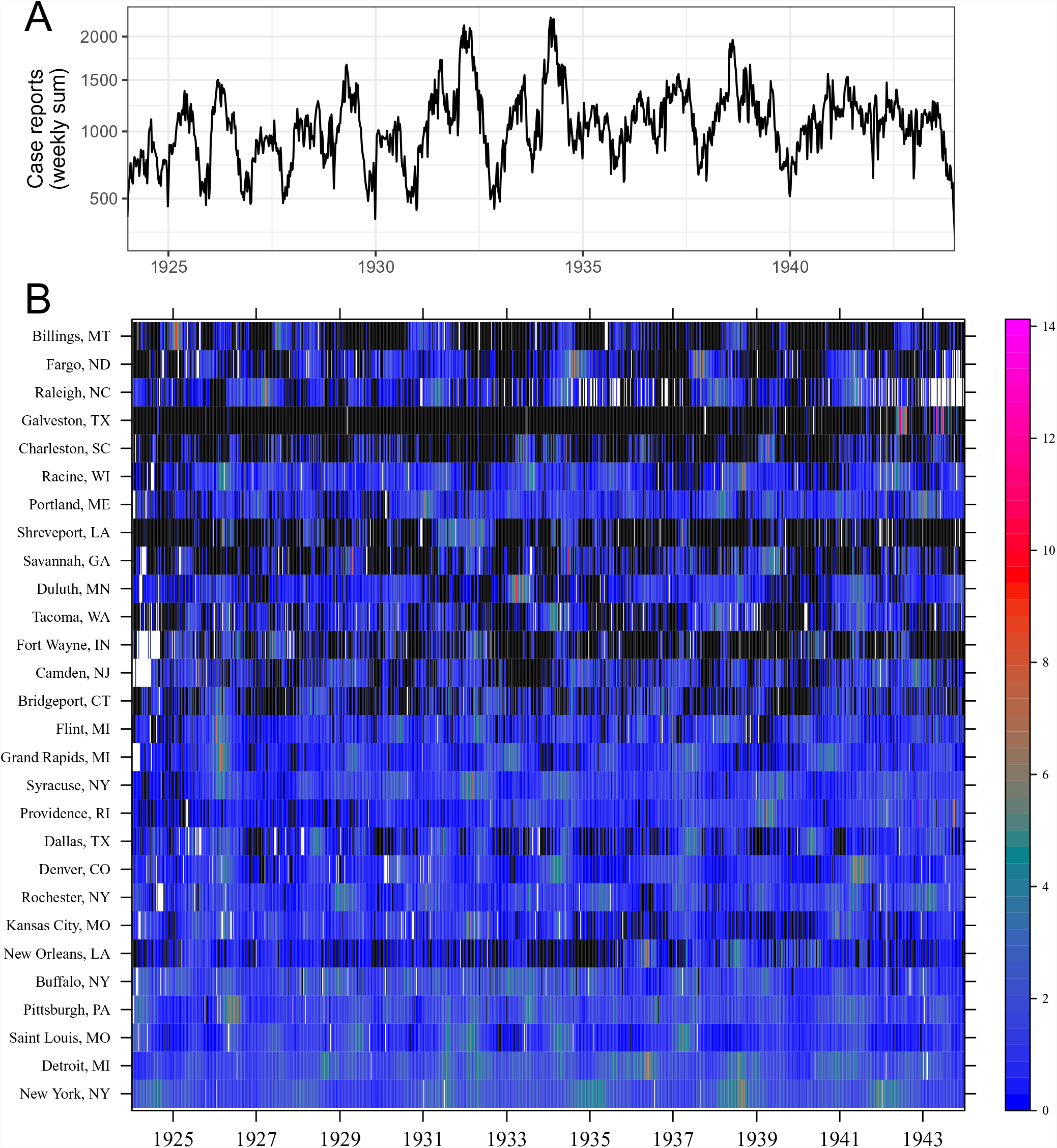
**Pertussis. A**. Total weekly case reports (all cities, sqrt-transformed). **B**. Weekly case reports (black = 0; white = missing). One city per row, ordered by population size, showing every third city. To facilitate comparison between cities, each city’s observations are scaled by that city’s temporal variance.

